# Neonatal hepatic myeloid progenitors expand and propagate liver inflammation in mice

**DOI:** 10.1101/2022.06.18.496674

**Authors:** Anas Alkhani, Cathrine Korsholm, Sarah Mohamedaly, Claire S. Levy, Caroline C. Duwaerts, Eric M. Pietras, Amar Nijagal

**Affiliations:** Department of Surgery, University of California, San Francisco, CA, USA; The Liver Center, University of California, San Francisco, CA, USA; Department of Comparative Pediatrics and Nutrition, University of Copenhagen, Denmark; Department of Medicine, University of California, San Francisco, CA, USA; Division of Hematology, University of Colorado Anschutz Medical Campus, CO, USA; The Pediatric Liver Center at UCSF Benioff Childrens’ Hospital, San Francisco, CA, USA; Eli and Edythe Broad Center of Regeneration Medicine, University of California, CA, USA

**Author notes:** **Correspondence:** Amar Nijagal, MD, Assistant Professor of Surgery, Division of Pediatric Surgery, 513 Parnassus Avenue, HSW 1652, Campus Box 0570, University of CA, San Francisco, San Francisco, CA 94143-0570. AA and CK are co-first authors. **Disclosures:** The authors have no disclosures to report. **Data Transparency:** The original contributions presented in the study are included in the article/Supplementary Material; further inquiries can be directed to the corresponding author.

**Keywords:** biliary atresia, perinatal liver inflammation, hematopoietic stem and progenitor cells

## Abstract

**Background and Aims:** Biliary atresia is a rapidly progressive pediatric inflammatory disease of the liver that leads to cirrhosis and necessitates liver transplantation. The rapid progression from liver injury to fulminant liver failure in children with biliary atresia suggests that factors specific to the perinatal hepatic environment are important for disease propagation. Hematopoietic stem and progenitor cells (HSPCs) serve as central hubs of inflammation and rely on inflammatory signals for their emigration from the liver to the bone marrow in neonatal mice. We hypothesized that HSPCs are critical for the propagation of perinatal liver inflammation (PLI).

**Methods:** Newborn BALB/c mice were injected intraperitoneally with 1.5×10^6^ focus forming units of Rhesus Rotavirus (RRV) to induce PLI or with PBS as control. Livers from RRV- and PBS-injected mice were compared using histology and flow cytometry. To determine the effects of HSPCs on perinatal inflammation, RRV-infected neonatal mice were injected with anti-CD47 and anti-CD117 to deplete HSPCs.

**Results:** RRV-induced PLI led to a significant increase in the number of common myeloid progenitors (Flt3^**+**^ CMPs: PBS=4426±247.2 vs RRV=9856±2009, p=0.0316; Flt3^**-**^ CMPs: PBS=3063±254.9 vs RRV=9743±1539, p=0.0012). We corroborated these findings by observing a significant increase in CD34^+^ hematopoietic progenitors/cm^2^ in histological sections of RRV-infected livers (PBS=4.977±2.573 vs RRV=27.09±12.49, p=0.0075). Elimination of progenitors through antibody-mediated myeloablation rescued animals from PLI and significantly increased survival (RRV+isotype control 55.56% vs RRV+myeloablation 94.12%, Chi-test=0.01).

**Conclusions:** These data demonstrate that RRV causes expansion of HSPCs and propagates PLI. Targeting of HSPCs may be useful in preventing and treating neonatal inflammatory diseases of the liver like biliary atresia.

**SYNOPSIS:** Hematopoietic progenitors reside in juvenile mouse livers even after the main site of hematopoiesis has shifted to the bone marrow. These progenitors are critical for the pathogenesis of perinatal liver inflammation as myeloablation rescues animals from disease.

## INTRODUCTION

Biliary atresia (BA) is the leading cause of pediatric liver transplants worldwide[1]. Though its exact etiology is unknown, the progressive perinatal liver inflammation (PLI) observed in patients with BA is caused by dysregulated immune responses to liver injury[2, 3]. Our recent studies in mice indicate that the adaptive immune response plays a limited role in the pathogenesis of PLI, whereas myeloid populations are critical for determining disease outcome as the relative proportions of pro-inflammatory and pro-reparative myeloid cells control disease severity[4]. The rapid progression from liver injury to fulminant liver failure in children with BA suggests that factors specific to the fetal hepatic environment are important for disease propagation. Therefore, understanding the fetal hepatic immune environment in which perinatal inflammatory diseases like BA develop and progress is important to identify promising treatment strategies for this devastating disease.

Hematopoietic stem and progenitor cells (HSPC) give rise to immune populations and reside in the liver of late-gestation human fetuses before emigrating to the bone marrow (BM)[5, 6]; in mice, this transition occurs during the first weeks of postnatal life [7]. HSPCs react to inflammatory signals and act as central hubs to coordinate immune responses, and through their rapid expansion and replenishment of mature myeloid populations, HSPCs are fundamental in facilitating the transition from inflammation to resolution of liver disease[4, 8-10]. For example, both human and murine models of liver injury show that monocytes transition from pro-inflammatory cells immediately after injury into pro-reparative monocytes once the injurious agent is no longer present[4, 9-11]. Our group has previously elaborated on this observation by demonstrating that an abundance of pro-reparative Ly6C^Lo^ non-classical monocytes render animals resistant to PLI, and that when the number of Ly6C^Lo^ non-classical monocytes is reduced, susceptibility to liver injury is restored[4]. In addition to the rapid expansion and differentiation of HSPCs into myeloid cells that occurs during inflammation, dysregulation of HSPCs can contribute to a feed-forward loop that leads to the pathologic expansion of inflammatory myeloid cells, resulting in chronic inflammation[8]. Taken together, these findings support the role of HSPCs and their mature myeloid progeny in propagating PLI.

In this study, we hypothesized that HSPCs propagate PLI in neonatal mice. To test this hypothesis, we used an infectious model of PLI to examine the role of HSPCs in neonatal liver injury. Using this model, we compared HSPCs and mature myeloid populations from liver and BM in the setting of homeostasis and PLI. Our results demonstrate that HSPCs expand during PLI and that depletion of HSPCs prevents liver inflammation. These findings support our hypothesis that HSPCs play an important role in propagating PLI.

## RESULTS

### The liver is a reservoir for hematopoietic progenitors in neonatal mice

To define the distribution of myeloid progenitors in neonatal animals under normal conditions, we quantified HSPCs (HSC^LT^s, CMPs, TMPs) and their mature progeny in the liver and BM of neonatal (postnatal day 3, P3) and juvenile (postnatal day 14, P14) mice. All HSPC populations were identified using cell-surface markers: HSC^LT^ were defined as Sca-1^+^, and CMP (common myeloid progenitor) and TMP (terminal myeloid progenitor) populations were Sca-1^-^. Individual CMPs and TMPs were distinguished based on the expression of CD34, Fc1R, Flt3, Ly6C, and CD115 (**Figure 1a,b**)[12]. Since our previous work demonstrated a limited role for T and B lymphocytes in the pathogenesis of PLI[4], lymphoid progenitors were not quantified in this study. We first quantified the lineage-negative (Lin^-ve^) compartment in the liver and found that the percentage and number of Lin^-ve^ cells did not differ significantly between P3 and P14 liver (**Figure 1c**) and the number of specific HSPC populations, including CMPs, and TMPs, were significantly lower at P14 than at P3 (**Figure 1c,d**). These findings are consistent with the known emigration of HSPCs from the liver during the first 2-3 weeks of postnatal life in mice.

**Figure 1:**
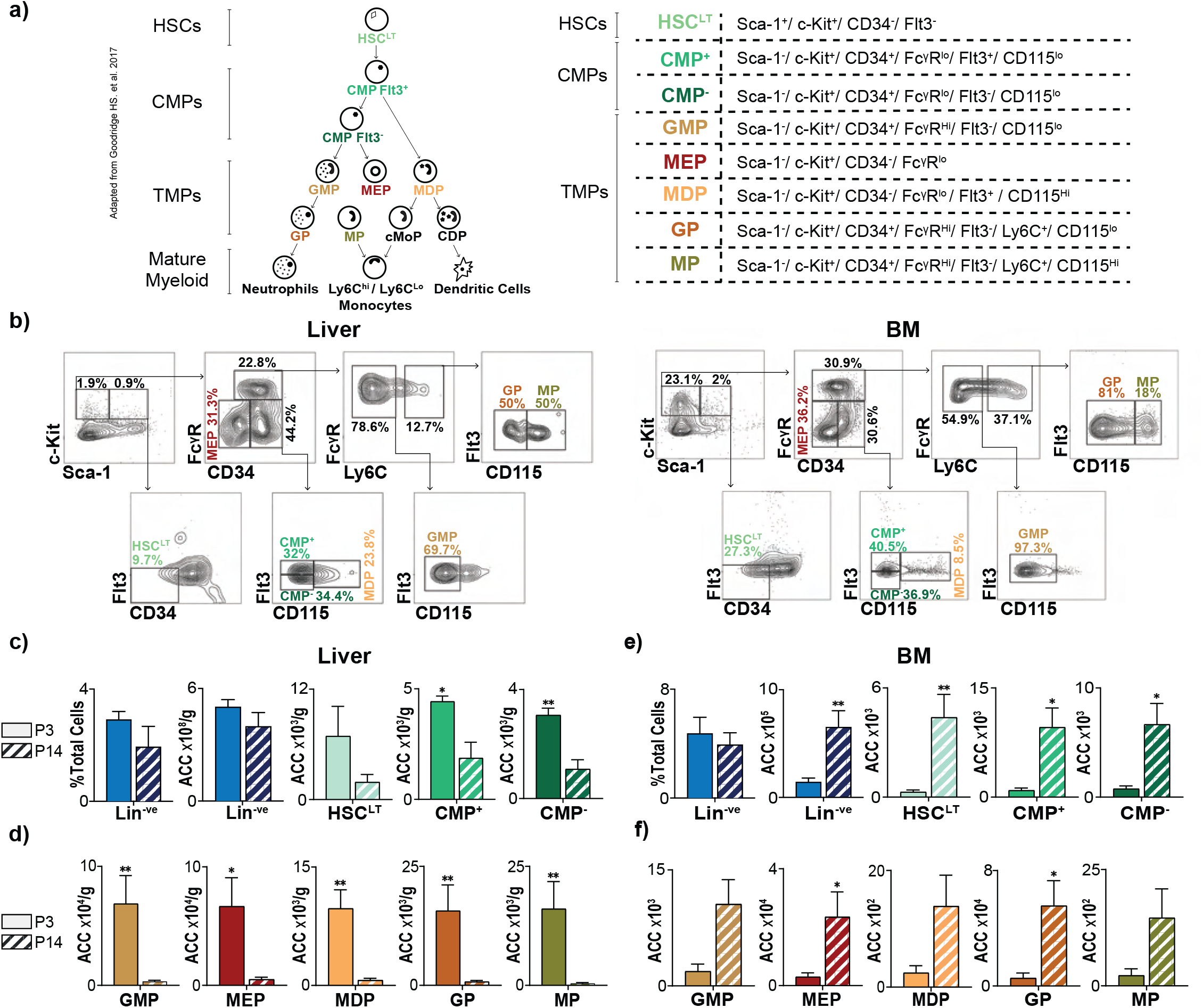
The liver is the main reservoir for myeloid progenitors in neonatal mice. **(a)** Schematic showing differentiation hierarchy and cell-surface markers of hematopoietic stem and progenitor cells (HSPCs): long-term hematopoietic stem cells (HSC^LT^), common myeloid progenitors (CMP^+^ and CMP^-^), and terminal myeloid progenitors (TMP: megakaryocytic-erythroid progenitors, MEP; monocytic-dendritic progenitors, MDP; granulocytic-monocytic progenitors, GMP; granulocytic progenitors, GP; monocytic and common monocytic progenitors, MP), and mature myeloid populations. **(b)** Plots demonstrating flow cytometric gating strategy of HSC^LT^s, CMPs, and TMPs in liver and bone marrow (BM) of PBS-injected mice. Quantification of lineage negative (Lin^-ve^) fraction and absolute cell counts (ACC) of Lin^-ve^, HSC^LT^, and CMP populations on post-natal day 3 (P3) and 14 (P14) of life in **(c)** liver and **(e)** BM. Quantification of TMPs in P3 and P14 **(d)** liver and **(f)** BM. Experiments are representative of n=6 for each group. p-value *<0.05; **<0.01. Error bars represent mean ± SEM.

In contrast to these findings in the liver, both the number of Lin^-ve^ cells and downstream progenitor populations (HSC^LT^, CMP^+^, CMP^-^, MEP, and GPs) in BM increased significantly between P3 and P14 (**Figure 1e,f**). In both the liver and BM, the mature myeloid populations mirrored the trends seen among progenitor populations as mature populations decreased in the liver and increased in the BM from P3 to P14; these differences, however, were not statistically significant (**Supplementary Figure 1a,b**). Collectively, these findings quantify the extent to which the murine liver retains hematopoietic progenitors during early neonatal life, and are consistent with the known migration of hematopoietic progenitors from the liver to the BM.

### The juvenile liver retains a higher proportion of common myeloid progenitors than do neonatal mice and maintains the ability to generate pro-myeloid colony forming units

We next asked whether the relative proportions of individual HSPC populations in the liver and BM changed from P3 to P14. In the liver, both the percentages of CMPs as a fraction of all Lin^-ve^ cells (**Figure 2a**) and as a fraction of HSC^LT^s, CMPs, and TMPs (**Figure 2b**) significantly increased between P3 and P14. Meanwhile, all liver TMPs decreased, although this was statistically significant only for MDPs and MPs (**Figure 2a,b**). Unlike the liver, the BM exhibited a relative increase in the percentages of HSC^LT^s, CMPs, and TMPs as a fraction of total Lin^-ve^ cells between P3 and P14 (**Figure 2c**). This increase was only significant for HSC^LT^, MEPs, and GPs, and not for CMPs. The same trend was observed when we examined each population as a fraction of all HSC^LT^s, CMPs, and TMPs (**Figure 2d**). These results highlight that the CMPs in the liver increase relative to other Lin^-ve^ cells, at a time when the main site of hematopoiesis is shifting to the BM.

**Figure 2:**
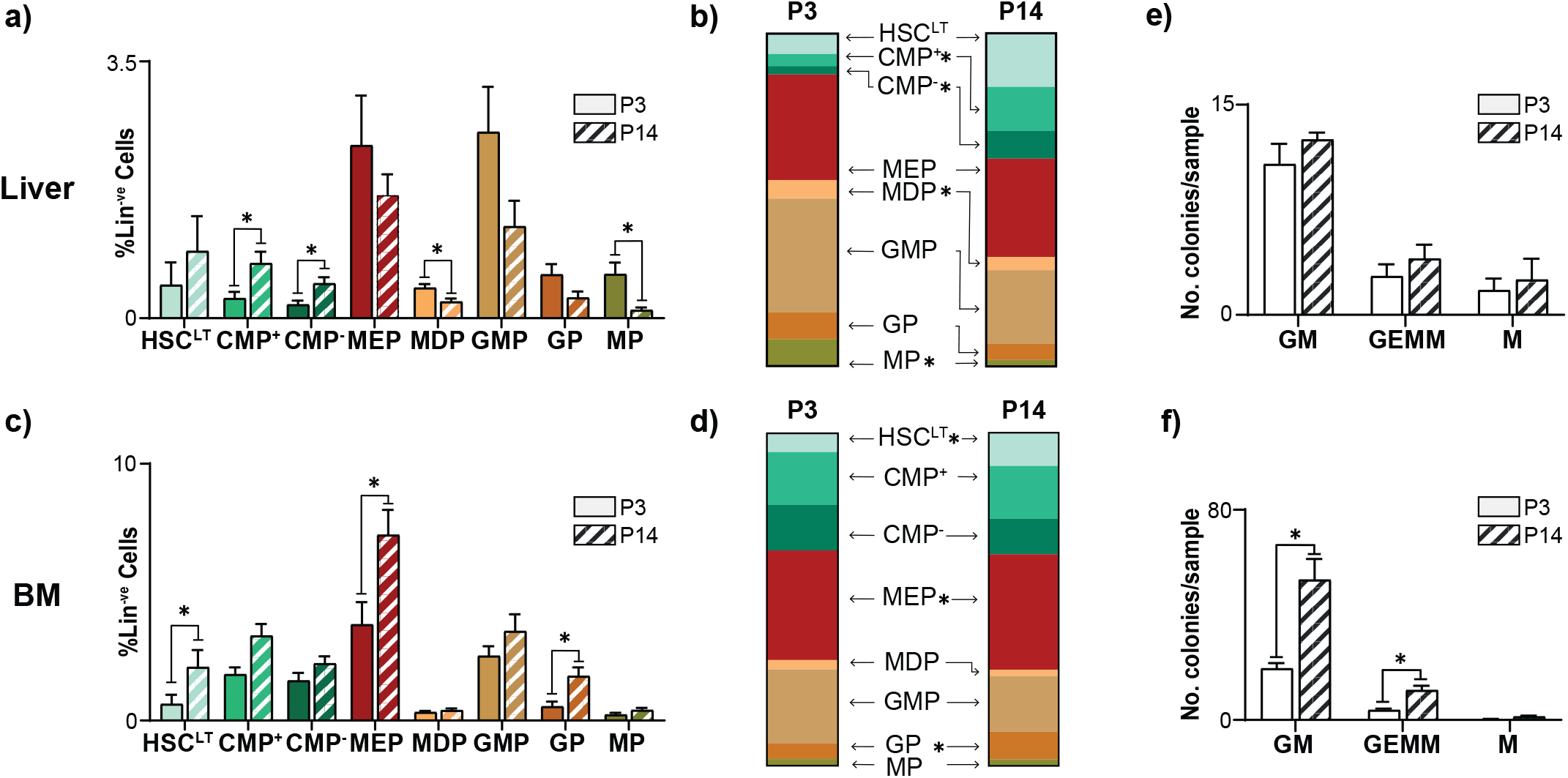
The juvenile mouse liver retains common myeloid progenitors and maintains myeloid differentiation capacity. Percentage of hematopoietic stem cell (HSC^LT^), common myeloid progenitor (CMP), and terminal myeloid progenitor (TMP) populations among lineage negative (Lin^-ve^) cells in **(a)** liver and **(c)** bone marrow (BM) on post-natal day 3 (P3) and 14 (P14). Relative proportions of HSC^LT^, CMP, and TMP among HSC^LT^ and their downstream myeloid progenitors in the **(b)** liver and **(d)** BM (P3, P14, n=6). Number (no.) myeloid colony forming units from **(e)** liver and **(f)** BM at P3 (liver n=8, BM n=5) and P14 (liver n=2, BM n=3). p-value *<0.05. Error bars represent the mean ± SEM.

To test the differentiation potential of HSPCs from the liver and BM, we quantified colony forming units (CFU) of pro-myeloid colonies. The liver retained a similar pro-myeloid differentiation capacity at P14 compared to P3 as the number of granulocyte monocyte (GM) and granulocyte, erythrocyte, monocyte, megakaryocyte (GEMM) and megakaryocytes (M) colonies remain unchanged (**Figure 2e**). In P14 BM, however, colonies from GM and GEMM increased, while the number of M colonies remained constant compared to P3 (**Figure 2f**). The increase in CMPs in the livers of juvenile mice and the maintenance of myeloid differentiation capacity led us to question whether HSPCs residing in the liver play a role in local immune responses to injury and inflammation.

### PLI leads to expansion of CMPs in the neonatal liver

Based on our observation that the liver is a reservoir for HSPCs in neonatal mice and the known role of HSPC populations as central hubs of inflammation, we questioned how PLI affects HSPCs. We have used rhesus rotavirus infection in neonatal mice to study the role of immune populations during PLI[4]. Neonatal pups injected with RRV within 24 hours of life develop progressive liver inflammation, and replicate the periportal inflammatory infiltrate seen in human BA[13]. To evaluate the effects of PLI, we analyzed the livers of RRV-injected pups using flow cytometry and histology three days after injury (P3). All HSPCs were identified from flow plots as shown in **Figure 3a**. Mature populations were identified using a previously published gating strategy[4]. PLI significantly increased the number of Lin^-ve^ cells in the liver (**Figure 3b**), which was reflected in downstream HSC^LT^s and CMPs (**Figure 3c**). PLI had no effect among TMPs (**Figure 3c**). When we assessed each progenitor population as a fraction of total HSC^LT^s, CMPs, and TMPs, we found that PLI led to increases in liver HSC^LT^ and CMP fractions (**Figure 3d**). Using immunohistochemistry to localize CD34^+^ HSPCs in the P3 liver, we found that PLI significantly increased the number of CD34^+^ HSPCs/cm^2^ relative to controls and these cells infiltrated all parts of the liver tissue with no identifiable pattern (**Figure 3e,f)**. Notably, the expansion of CMPs did not lead to an increase in mature myeloid populations at P3 (**Figure 3g,h**)

**Figure 3:**
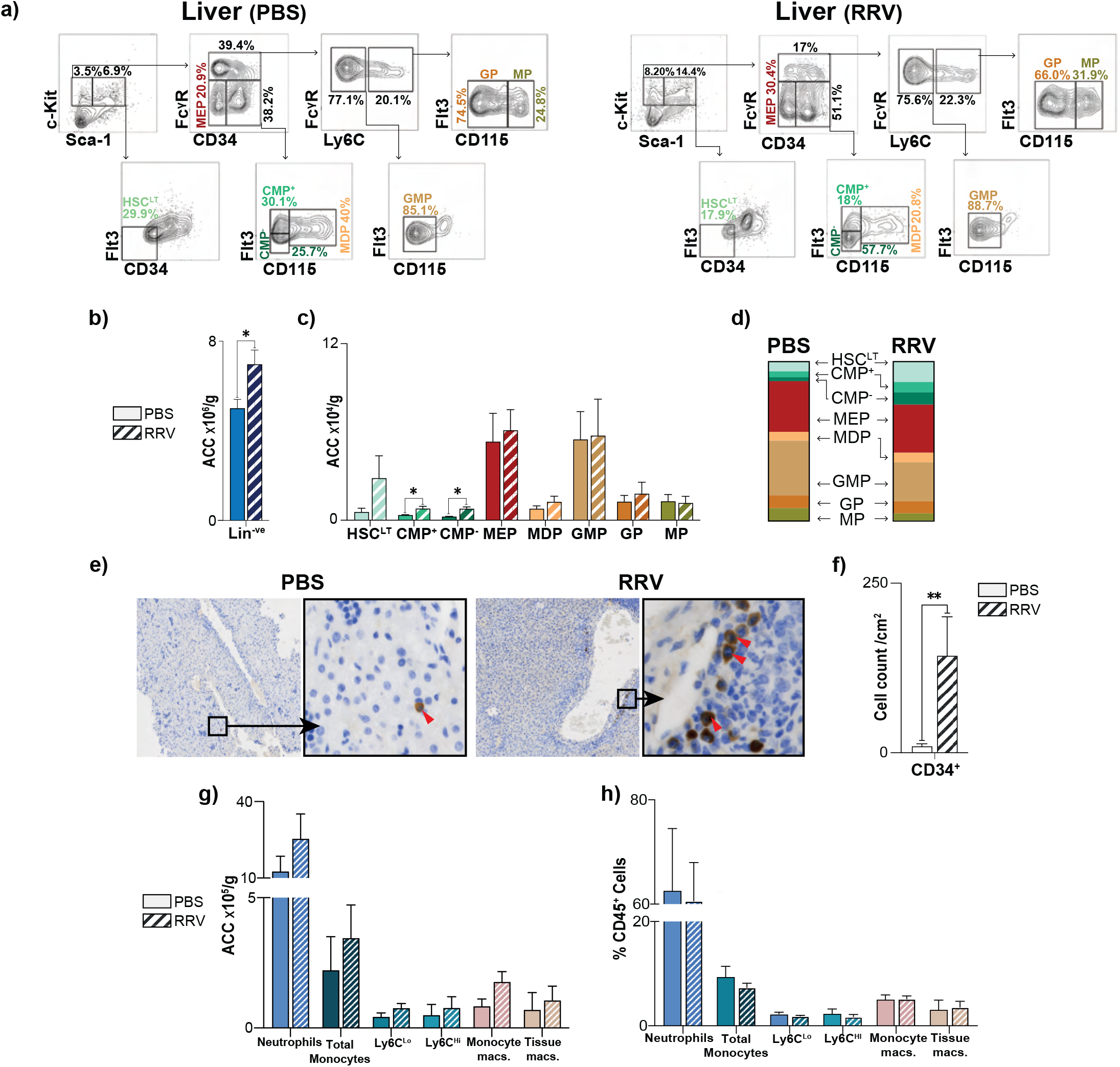
Perinatal liver injury results in the expansion of myeloid progenitors within the neonatal liver. **(a)** Representative flow plots demonstrating gating strategy of hematopoietic stem cells (HSC^LT^), common myeloid progenitors (CMP), and terminal myeloid progenitors (TMP) in liver of PBS- and RRV-injected 3-day-old (P3) mice. Quantification of absolute cell counts (ACC) of **(b)** lineage negative (Lin^-ve^) cells and **(c)** HSC^LT^s, CMPs, TMPs in P3 livers from PBS- (n=6) and RRV-injected (n=6) mice. **(d)** Relative fractions of HSC^LT^, CMP, and TMP populations in P3 livers from PBS- (n=6) and RRV-injected (n=6) mice. **(e)** Representative P3 livers in the setting of PBS and RRV with red arrows marking immunohistochemistry-stained CD34^+^ cells. **(f)** Quantification of CD34^+^ cells at P3 in PBS- (n=6) and RRV-injected mice (n=9). Quantification **(g)** ACC and **(h)** %CD45^+^ leukocytes of mature myeloid populations in the liver of PBS- (n=6) and RRV-injected (n=7) mice at P3. p-value *<0.05, **<0.01. Error bars represent mean ± SEM.

### PLI leads to contraction of HSPCs and expansion of mature myeloid populations in juvenile mice

We next sought to determine whether the expansion of CMPs seen in neonatal livers after RRV infection persisted or was a temporary increase during PLI. For this, we quantified HSPC populations 14 days after RRV infection in juvenile mice (P14) and observed a reduction of Lin^-ve^ cells **(Figure 4a)**. The number of HSPCs, including HSC^LT^, CMP, MEP, and MPs, in RRV-infected P14 livers decreased, compared to PBS injected controls. Despite the decrease in the number of these progenitor populations, only MEPs decreased as a proportion of Lin^-ve^ cells and as a fraction of all HSC^LT^s, CMPs, and TMPs **(Figure 4b,c)**. The percentage of neutrophils, monocytes, and monocyte-derived macrophages increased in livers of RRV-infected juvenile mice, corresponding to the peak of disease in RRV-infected mice **(Figure 4d,e)**[13]. These findings demonstrate that PLI causes a temporary expansion of CMPs in RRV-infected neonatal mice and results in a relative increase in mature myeloid populations 14 days after RRV infection.

**Figure 4:**
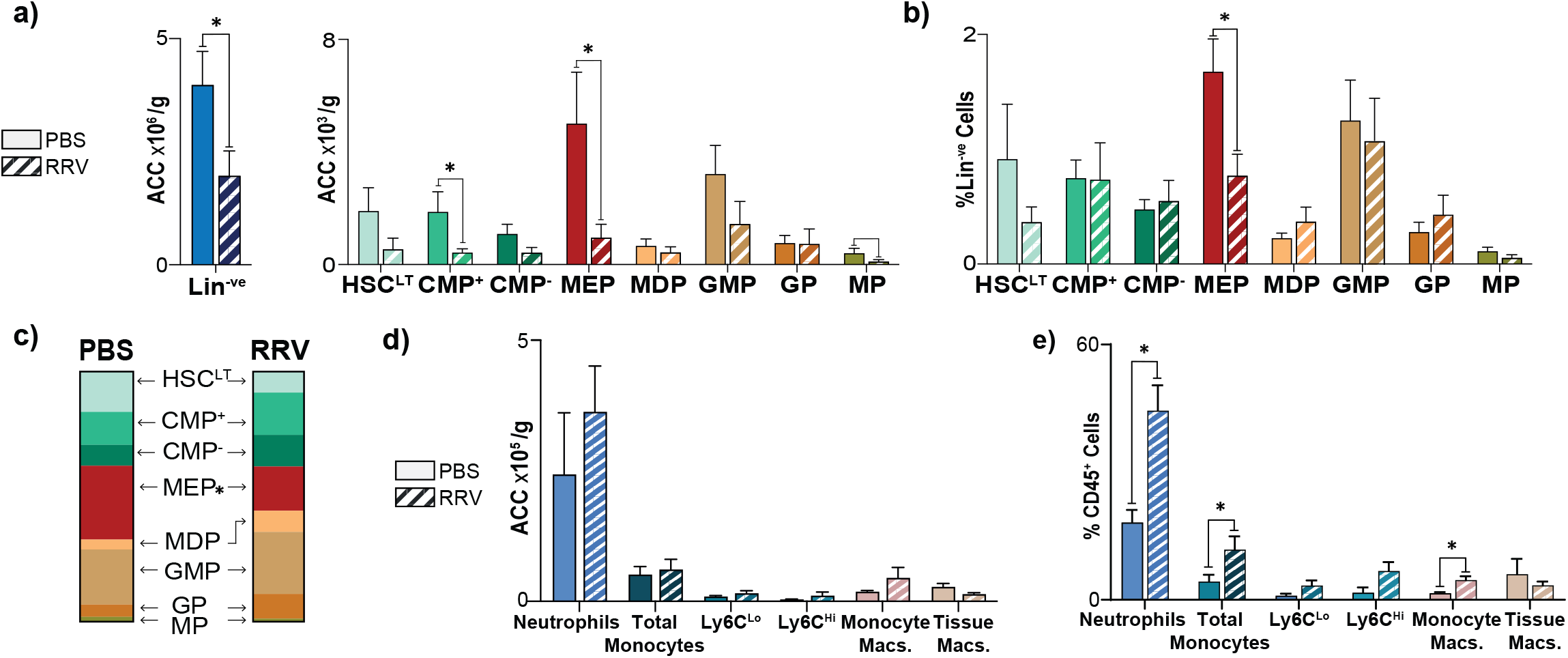
Perinatal liver inflammation leads to contraction of hematopoietic progenitors and expansion of mature myeloid cell proportions within the juvenile liver. **(a)** Quantification of lineage negative (Lin^-ve^) cells, hematopoietic stem cells (HSC^LT^), common myeloid progenitors (CMPs), and terminal myeloid progenitors (TMPs) of the liver in terms of absolute cell counts (ACC), **(b)** percentage of total Lin^-ve^ cells, **(c)** fraction of whole for HSC^LT^, CMPs, and TMPs. Quantification of mature myeloid populations (PBS n=3, RRV n=3) of the liver in terms of **(d)** ACC and **(e)** percentage of individual mature myeloid populations out of all CD45^+^ cells (PBS n=3, RRV n=3) at post-natal day 14 (P14). Experiments are representative of n=6 for PBS and n=8 for RRV unless otherwise stated. p-value *<0.05. Error bars represent mean ± SEM.

### Myeloablation protects mice from RRV-mediated perinatal liver inflammation

The expansion of CMPs in the neonatal liver after RRV infection led us to question whether HSPCs play a role in propagating PLI. To test this, we evaluated the effect of depleting HSPCs on the progression of PLI, using synergistic, myeloablating anti-CD117 and anti-CD47 antibodies[14, 15]. Mice were injected with RRV on day 0 after birth to induce PLI. From day 0-4, they were also injected with anti-CD117 + anti-CD47 (MA) or IgG2b + Ig2a isotype controls (Iso) (**Figure 5a**). In the MA group, 94% of the pups survived RRV compared to only 56% in the isotype control group. (**Figure 5b**). MA pups also weighed more than isotype controls, but this difference was not statistically significant (**Figure 5c**). The improvement in survival and weight of the MA pups was corroborated by fewer moribund features, such as hair loss, dehydration, hunched appearance, and jaundice (**Figure 5d**). Finally, the extent of liver injury significantly decreased in the MA pups, as evidenced by less periportal inflammation and no identifiable areas of hepatic necrosis (**Figure 5e**).

**Figure 5:**
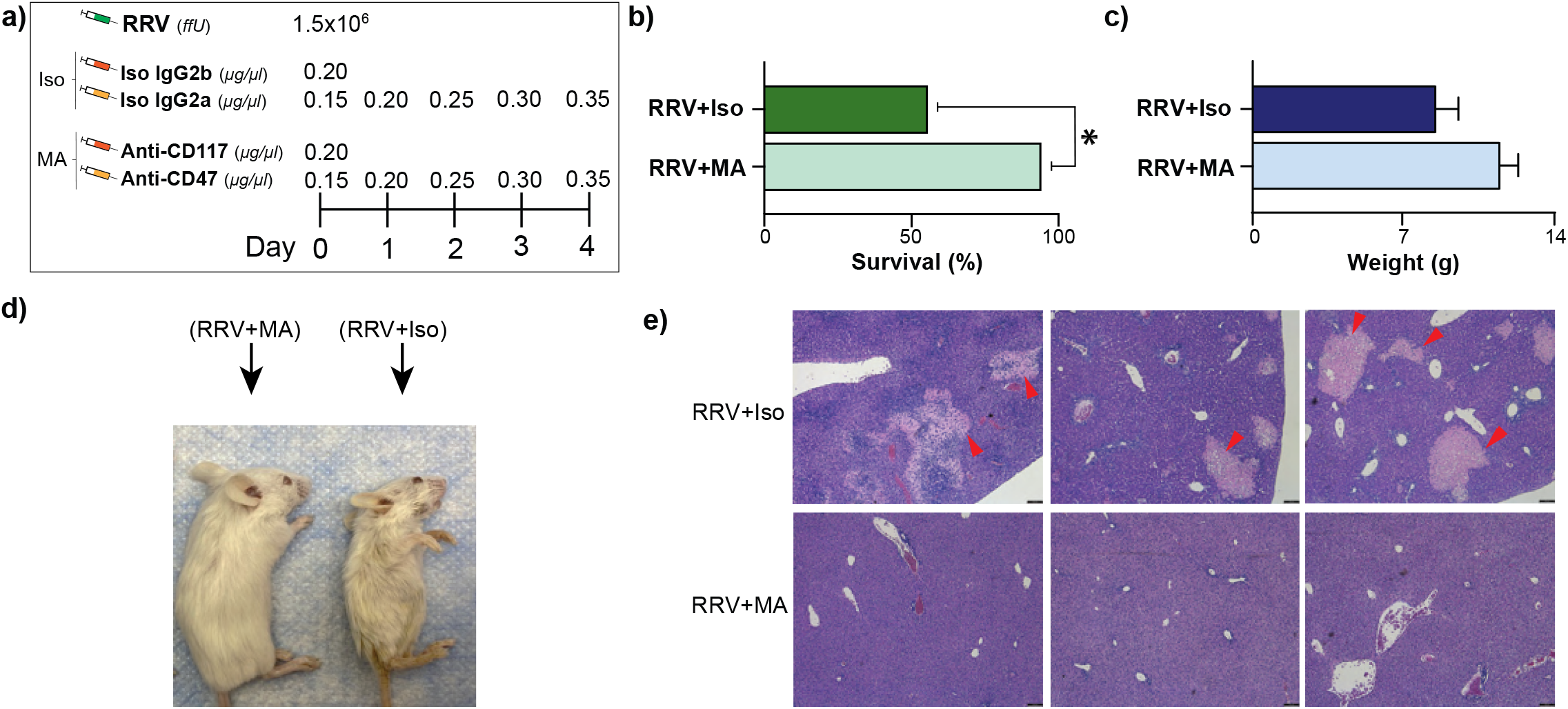
Antibody-mediated myeloablation protects mice from perinatal liver injury. **(a)** Dosage schedule illustrating Rhesus Rotavirus (RRV) injection, isotype (Iso)-injections, and the myeloablative (MA) injections used to deplete hematopoietic stem and progenitor cells (HSPCs). (b) Percent survival and **(c)** pup weights from mice three weeks after injection with either RRV+MA or RRV+Iso. Pictures illustrating **(d)** mice after RRV+MA and RRV+Iso, and **(e)** histological H&E sections of livers from animals in both groups. Red arrows points to the necrotic foci. Experiments are representative of n=17 for RRV+MA and n=9 for RRV+Iso. p-value *<0.05. Error bars represent mean ± SEM.

These results demonstrate that PLI not only causes expansion of HSPCs but also that the propagation of injury after RRV infection is dependent on HSPCs.

## DISCUSSION

In this study, we defined the composition of HSPCs during homeostasis in neonatal and juvenile mouse liver, and we used a mouse model of perinatal liver injury to define the changes that occur to HSPCs during PLI. We found that 1) common myeloid progenitors reside in the livers of juvenile mice even after the main site of hematopoiesis has shifted to the BM, 2) PLI leads to the expansion of common myeloid progenitors in the neonatal liver and causes an expansion of mature myeloid progenitors in juvenile animals, and 3) targeted depletion of HSPCs using anti-CD117 and anti-CD47 prevents RRV-induced PLI, as demonstrated by improved survival, jaundice clearance, and decreased hepatic necrosis.

Our results demonstrate that under homeostatic conditions, the neonatal mouse liver continues to act as a reservoir for common myeloid progenitors (CMPs). Furthermore, the differentiation capacity of HSPCs towards the myeloid lineage persists in the liver even after the main hematopoietic site is shifting to the BM. These findings corroborate the known role of the adult liver as an important immunogenic organ and the first-line of defense against pathogens of the gut[11], such as RRV[16]. Though recruitment of myeloid cells (neutrophils, monocytes) during inflammation has been thought to occur from the blood stream[11], our results supports the idea that remnant HSPCs in the liver may serve as central hubs of inflammation during PLI.

Our findings also support the idea that liver inflammation during perinatal life affects emigrating hepatic progenitor populations. In humans, the spatial and temporal overlap of liver development and hematopoiesis in the late-gestational fetus[17] may contribute to the devastating acute and chronic consequences of both liver- and immune function observed in progressive inflammatory diseases like BA. In our study, perinatal liver injury led to an early increase in HSPCs and depletion of these cells lessened clinical and histological signs of liver inflammation. Similar findings of HSPCs driving continuous inflammation have previously been observed in the heart, where chronic inflammation transforms HSPCs into a pro-inflammatory phenotype that enhances inflammation in a destructive feed-forward loop[8]. Our findings indicate that this detrimental feed-forward loop is similarly present in BA, where an injury to the fetal liver leads to dysregulation of HSPCs and propagation of inflammation.

The current treatment of BA relies on early surgical treatment to restore bile flow after Kasai portoenterostomy, although most patients will continue to develop progressive liver injury requiring liver transplantation, highlighting the need for new and innovative treatments. Our results indicate that HSPCs propagate perinatal liver injury in mice and suggest that HSPCs may also contribute to human BA. Intriguingly, manipulation of HSPC populations have been found to influence disease outcome in human patients. In infants with BA after Kasai portoenterostomy, the effect of administering three consecutive days of granulocyte-colony-stimulating-factor (G-CSF) on liver inflammation was examined in a phase 1 trial showing that peripheral neutrophils and HSPCs initially increased before decreasing to baseline levels after two weeks. Notably, G-CSF treatment was associated with reduced cholestasis at one month after treatment began but reverted to control levels after three months[18]. A phase 2 randomized controlled trial is currently underway to determine the efficacy of G-CSF in patients with BA (NCT04373941). Our findings and those from the phase I trial highlight that HSPC populations are dynamic during the course of an inflammatory insult and their functions change depending on the stage of disease and age of the patient. These findings also support the idea that manipulation of specific HSPC subsets may prove to be efficacious in resolving BA.

The changes to HSPCs after RRV infection support the idea that early perinatal inflammatory insults have long-term consequences to immune function. Children with BA, for example, have an increased infection rate and decreased vaccine responses compared to healthy controls[19, 20] and they are more likely than children with diseases other than BA who received liver transplants to reject donor livers[21]. In mice, a primary pathogenic stimulus has been found to cause changes in epigenetic and translational properties of immune cells that leads to a heightened response to similar stimuli in the future – a concept known as ‘trained immunity’[22, 23]. As myeloid cells account for the primary immune response during PLI[4], trained immunity may play a role in the long-term immune dysregulation observed in BA.

In conclusion, our study demonstrates that myeloid progenitors increase during PLI and that their depletion improves disease outcome. Future studies are necessary to investigate the specific effects of myeloablation on myeloid progenitor populations in neonates. Our study does suggest, however, that targeting hematopoietic progenitors may be useful in preventing and treating neonatal liver inflammatory disease like BA.

## METHODS

### Mice

Balb/C mice were obtained from the National Cancer Institute (Wilmington, MA, USA), and were cared for according to the *Guide for Care and Use of Laboratory Animals*. Mouse experiments were approved by University of California, San Francisco Institutional Animal Care and Use Committee, and all mice were euthanized according to humane endpoints.

### Creation of single-cell suspensions

All livers were isolated and mechanically dissociated in phosphate buffered saline (PBS). Livers from day P14 mice were enzymatically digested using 2.5 mg/mL liberase (Roche Indianapolis, IN, 05401119001) in 1 M CaCl_2_ HEPES buffer. BM was isolated from the tibia, fibula, hip, and lower spines of neonatal animals, and mechanically dissociated in PBS. Both liver and BM single cell suspensions were filtered through a 100um strainer prior to further analysis.

### Flow Cytometry

In order to analyze HSPCs, single-cell suspensions from liver and BM were depleted of lineage-positive (Lin^+^) cells using a Direct Lineage Cell Depletion Kit (Miltenyi Biotec, 130-110-470) and stained for the following HSPCs: long-term hematopoietic stem cells (HSC^LT^), common myeloid progenitors (CMPs), and terminal myeloid progenitors (TMPs) using cell surface markers (Supplementary Tables 1 and 2). Flow cytometry was performed on a LSRFortessa X20 (BD Biosciences) and data were analyzed in FlowJo (Ashland, OR).

### Colony-forming unit (CFU) assays

Single-cell suspensions from day P3 and P14 livers and BMs were cultured on MethoCult− media containing methylcellulose, recombinant mouse stem cell factor, IL-3, IL-6, and recombinant human EPO. 2×10^4^ cells were plated and incubated at 37°C in 5% CO_2_ for 12 days. The proliferation and differentiation ability of HSPCs was assessed by a blinded observer who categorized the following colonies: granulocyte, macrophage (GM), granulocyte, erythrocyte, macrophage, megakaryocyte (GEMM), and macrophage (M).

### Postnatal model of perinatal liver inflammation

Rotavirus (RRV) was grown and titered in *Cercopithecus aethiops* kidney epithelial (MA104) cells. PLI was induced by intraperitoneal injections (i.p.) of 1.5×10^6^ focus forming units (ffu) RRV within 24 hours of birth (P0). Controls were injected i.p. with PBS.

### Histologic analysis

Liver tissue was stained with immunohistochemistry (IHC) for CD34^+^ cells or hematoxylin and eosin (H&E). IHC slides were imaged at 40x magnification and CD34^+^ cells with large nucleus and little cytoplasm were counted as HSPCs by a blinded observer using QuPath[24]. Since CD34 is also present on vascular endothelial cells, all elongated cells with the morphologic appearance of endothelial cells were excluded[25]. CD34^+^ cells/cm^2^ were calculated within an entire tissue section using QuPath[24].

### Antibody-mediated myeloablation in neonatal mice

Myeloablation was induced by i.p. injections (20ul) of anti-CD117 and anti-CD47. Anti-CD117 (0.20 ug/ul) was given only on day 0. Anti-CD47 was given on day 0 (0.15 ug/ul), day 1 (0.20 ug/ul), day 2 (0.25 ug/ul), day 3 (0.30 ug/ul), and day 4 (0.35 ug/ul) post RRV-injection. Isotype controls were injected i.p. with isotype IgG2b (similar regimen as anti-CD117) and isotype IgG2a (similar regimen as anti-CD47). Escalating amounts of anto-CD47 and IgG2a were given to account for the natural increase in pup weight that occurs after birth. All antibodies for this experiment were purchased from BioXCell, West Lebanon, NH.

## Abbreviations

ACC: Absolute cell count
BA: Biliary Atresia
BM: Bone Marrow
CFU: Colony forming unity
CMP: Common myeloid progenitors
ffu: Focus forming units
G-CSF: Granulocyte-colony-stimulating-factor
GEMM: Granulocyte, erythrocyte, macrophage, megakaryocyte
GM: Granulocyte, macrophage
GMP: Granulocytic-monocytic progenitors
GP: Granulocytic progenitors
HSC^LT^: Long-term Hematopoietic Stem Cells
MA: Myeloablation
MDP: Monocytic-dendritic progenitors
MEP: Megakaryocytic-erythroid progenitors
MP: Common monocytic progenitors
PLI: Perinatal liver inflammation
RRV: Rhesus rotavirus
TMP: Terminal Myeloid Progenitors

## Data analysis

All graphs and statistics were generated using GraphPad Prism 9.3.1 (San Diego, CA). Individual proportions of HSPCs were calculated, based on absolute cell counts (ACC), as either a percentage (%) of the total lineage-negative progenitor compartment (Lin^-ve^) cells, or as a fraction of all HSC^LT^ and downstream myeloid progenitors (CMPs and TMPs). Mature myeloid cell proportions were calculated as a percentage of CD45^+^ leukocytes based on ACC. p-values were calculated using unpaired, non-parametric (Mann-Whitney) tests except for survival calculations that were performed using chi-squared tests. A p-value of <0.05 was considered significant. Error bars represent mean ± standard error of the mean (SEM). All authors had access to the study data and had reviewed, and approved the final manuscript.

## Acknowledgements

The authors would like to thank Dr. Henry Greenberg (Stanford University, CA) for providing MA104 cells and Rhesus rotavirus. The authors would also like to acknowledge Dr. Scott Kogan for assistance with CFU assay interpretation, Dr. Kyle Cromer and Pamela Derish for their critical review of the manuscript. CK was supported by a fellowship from the Lundbeck Foundation’s Danish-American Research Exchange Program, administered by Innovation Center Denmark, Silicon Valley. Additional funding was provided by the American Pediatric Surgical Association Foundation Jay Grosfeld, MD Scholar Award (AN), an American College of Surgeons Faculty Research Fellowship (AN), a UCSF Liver Center Pilot Award (NIH P30 DK026743, AN), the UCSF Parnassus Flow Cytometry Core (DRC Center Grant NIH P30 DK063720), and core resources of the UCSF Liver Center (P30 DK026743).

