## Supplementary Data for "Neonatal hepatic myeloid progenitors expand and propagate liver inflammation in mice"

**SUPPLEMENTARY TABLES**

**Table 1: Staining panel for hematopoietic stem- and progenitor cells (HSPCs): Long-term hematopoietic stem cells ($HSC^{LT}$), common myeloid progenitors (CMPs), and terminal myeloid progenitors (TMPs)**

| Antibody | Clone | Manufacturer | Catalog number |
|---|---|---|---|
| c-kit CD117 | 2B8 | Biolegend, San Diego, CA | 105820 |
| Sca-1 Ly6A/E | D7 | Biolegend, San Diego, CA | 108114 |
| FcgR Cd16/Cd32 | 2.4G2 | BD Biosciences, Franklin Lakes, NJ | 553142 |
| LY6C | HK1.4 | Biolegend, San Diego, CA | 128012 |
| CSF-1R CD115 | AFS98 | Biolegend, San Diego, CA | 135506 |
| Flt3 CD135 | A2F10.1 | BD Biosciences, Franklin Lakes, NJ | 560718 |
| CD34 | RAM34 | BD Biosciences, Franklin Lakes, NJ | 553733 |

**Table 2: Staining panel for mature myeloid cells**

| Antibody | Clone | Manufacturer | Catalog number |
|---|---|---|---|
| MHCII | M5/114.15.2 | eBioscience, Waltham, MA | 48-5321-82 |
| Cd11b | M1/70 | eBioscience, Waltham, MA | 47-0112-82 |
| Cd45 | 30-F11 | eBioscience, Waltham, MA | 56-0451-82 |
| Ghost | *Not relevant* | Tonbo Biosciences, San Diego, CA | 13-0870-T100 |
| Ly6C | AL-21 | BD Biosciences, Franklin Lakes, NJ | 563011 |
| Ly6G | 1A8 | BD Biosciences, Franklin Lakes, NJ | 560601 |
| Fc Block Cd16/Cd32 | 2.4G2 | BD Biosciences, Franklin Lakes, NJ | 553142 |
| Cd11c | N418 | Biolegend, San Diego, CA | 117339 |
| Cd64 | X54-5/7.1 | Biolegend, San Diego, CA | 139323 |

**SUPPLEMENTARY FIGURE LEGENDS**

**Supplementary Figure 1:** *Mature myeloid populations contract in the liver while expanding in the bone marrow of the juvenile mouse.* Quantification of absolute cell count (ACC) of mature myeloid populations on day 3 (P3 n=3) and 14 (P14 n=3) in the **(a)** liver and **(b)** bone marrow (BM). Error bars represent mean ± SEM. Ly6C$^{Lo}$ non-classical monocytes (Ly6C$^{Lo}$), Ly6C$^{Hi}$ classical monocytes (Ly6C$^{Hi}$), Macrophages (macs)

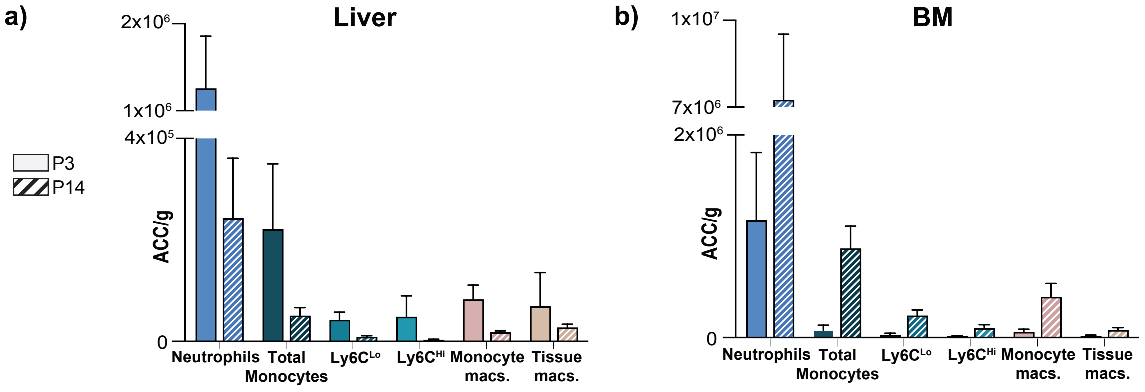


**Supplementary Figure 1:** *Mature myeloid populations contract in the liver while expanding in the bone marrow of the juvenile mouse.* Quantification of absolute cell count (ACC) of mature myeloid populations on day 3 (P3 n=3) and 14 (P14 n=3) in the **(a)** liver and **(b)** bone marrow (BM). Error bars represent mean ± SEM. Ly6C$^{Lo}$ non-classical monocytes (Ly6C$^{Lo}$), Ly6C$^{Hi}$ classical monocytes (Ly6C$^{Hi}$), Macrophages (macs)
